# Acquisition of Temporal Order Involves a Reverberating Network in Hippocampal Field CA3

**DOI:** 10.1101/623025

**Authors:** B.G. Gunn, B.M. Cox, C.D. Cox, A.A. Le, V.C. Inshishian, C.M. Gall, G. Lynch

## Abstract

Here we report that hippocampal field CA3 maintains self-sustained activity for remarkable periods following a brief input and that this effect is extremely sensitive to minor perturbations. Using novel behavioral tests, that as with human episodic learning do not involve training or explicit rewards, we show that partial silencing of the network in mice blocks acquisition of temporal order, but not the identity or location, of odors. These results suggest a solution to the question of how hippocampus adds time to episodic memories.

## Introduction

The manner in which the hippocampus adds the critical temporal dimension to an episodic memory is poorly understood. Learning theories generally posit that events closely associated in time become connected in memory^1^, an assumption that finds support in the temporal contiguity requirement (‘Hebb Rule’)^2^ for induction of long-term potentiation (LTP). How then are events separated by tens of seconds or longer linked together, as occurs routinely in episodic memory? Recent studies using prior training have identified multiple regions in the rodent hippocampus and entorhinal cortex in which the firing of neurons changes in a manner indicative of temporal processing^3-7^. These results describe potential substrates for the encoding of episodic elements but do not address the question of how the temporal order is acquired during first time, unsupervised sampling of a cue series. This operation is well recognized as being essential for the formation of human episodic memories and shown in recent studies on brain-damaged individuals to depend on the hippocampus^8^. A solution, first advanced decades ago, to the problem of bridging widely spaced cues postulates the existence of recurrent neuronal networks capable of maintaining reverberating (self-sustained) activity for extended periods ^1,9,10^. A number of investigators have suggested that the extremely dense commissural/associational (C/A) feedback system found in hippocampal field CA3^11^ constitutes such a system^1,9,10,12-14^ but experimental support is lacking.

We used hippocampal slices to conduct first tests for prolonged reverberating activity following activation of field CA3 and found that a brief input increased firing for a period (minutes) that is unprecedented for brain networks. Complex systems are prone to catastrophic collapse and this proved to be the case for self-sustained activity in CA3. This last result suggested an experimental design for testing if reverberation in the network plays an essential role in the acquisition of ‘when’ data. Using this, we found that suppression of the C/A system entirely blocks retention of the order in which cues had been sampled but not of their identities or locations.

## METHODS

### Animals

Studies used 2-6 mo old male mice (FVB-129 and C57BL6 backgrounds) group housed (3-5 per cage) with food and water ad libitum. Experiments were conducted in accord with NIH guidelines for the Care and Use of Laboratory Animals and protocols approved by the Institutional Animal Care and Use Committee.

### Behavioral Tests

Paradigms used olfactory cues to evaluate the ‘what’, ‘where’, and ‘when’ components of episodic-type memories in FVB-129 mice.

#### ‘What’

A serial odor task was used to assess encoding of cue identify (‘what’) information. Odorants were pipetted onto filter paper (final concentration of 0.1 Pascals) which was placed in a glass jar (5.25 cm diameter × 5 cm height) with a plastic lid containing a ∼1.5 cm diameter hole to allow for sampling. During the habituation session, the mouse was allowed to explore for 5 min a plexiglass test arena (30 × 25 cm floor, 21.5 cm walls) containing two cups (without odor). The mouse was then moved to an identical empty chamber for 5 min while the cups in the test arena were replaced with ones containing odor A (in duplicate) for the next session. After 3 min of exploration, the animal was again moved to the alternate chamber for 5 min and the cups with odor A in the test chamber were replaced with cups containing odor B. This sequence was repeated with odor C. In the final test session 5 min later the mouse was exposed to two different odors: a previously sampled odor A and a novel odor D. The time spent sampling D vs A was used as a measure of retention.

#### ‘Where’

This paradigm used a large plexiglass arena (60 cm × 60 cm floor, 30 cm walls) containing 4 odor cups. Mice were habituated to the chamber for 5 min with cups not containing odors and then transferred to an alternate identical empty holding chamber for 5 min. During the following training session the mouse was returned to the initial arena and allowed to explore cups scented with four different odors for 5 min. After placement in the holding chamber (5 min) the mouse was returned to the original arena in which the location of two of the four scented cups had been switched. In this final 5 min test session, the times spent exploring odors in the new and familiar locations were compared.

#### ‘When’

This paradigm was largely identical to the ‘what’ task, except that an additional odor pair D:D was added to the last step in the training sequence (see Fig 1c). After exposure to odor pair D, the final test session presented two different odors selected from the sequence A:A to D:D. Untreated mice explored the odor sampled least recently (e.g., odor B more than C).

**Figure 1.**
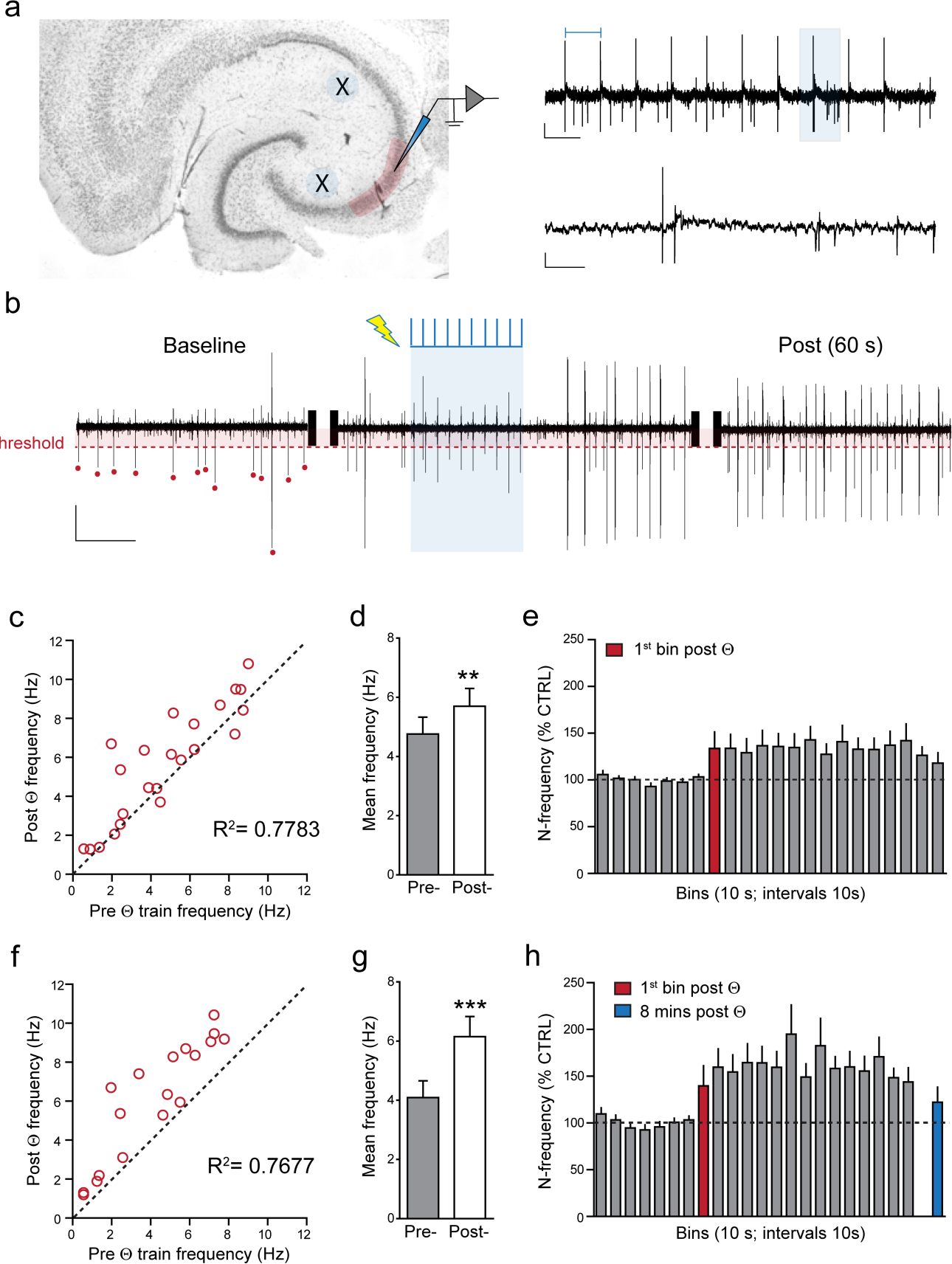
Brief physiological activation of the CA3 C/A system initiates autonomous cell firing lasting minutes. **a)** Nissl stained cross section of temporal hippocampus illustrates placement of stimulating (X) and recording (shaded) electrodes. Representative responses (right) during a 10-pulse Θ train with stimulation artifacts (top left bar), where an individual fEPSP (from shaded zone) is shown on an expanded time scale below (Scale bars: y=100 µV, x=200ms and 20ms for top and bottom traces, respectively). **b)** Representative sections of filtered (band pass 300-3000 Hz) extracellular recordings of PC spiking during the baseline period (4 s), the Θ stimulation train (shaded blue, 2 s) and subsequent 4 s, as well as 1 min afterwards (4 s). The detection threshold and those spikes counted during baseline are indicated with red shading and red dots, respectively. Note that firing frequency is increased following stimulation and that this is maintained for ≥1 minute (Scale bars: y=100 µV, x=1s). **c)** Scatter plot with unity line (dashed) illustrates the mean frequency before and 2 min after Θ stimulation for all cases (n=23 slices; 57 trials). **d)** Bar graph summarizing the mean frequency pre-and 2 min post stimulation for all cases (**p<0.003 paired t-test pre-vs. post). **e)** Bar graph summarizing the mean normalized (N) frequency (% control, CTRL) for each 10 s bin during the baseline period and for 4 min after Θ stimulation for 23 slices: group effects for bins were significant (p<0.0001 pre-vs post stimulation; RM-ANOVA). **f)** Scatter plot with unity line (dashed) summarizes the mean frequency before and for the 2 min after Θ stimulation for positive cases only (n=19 slices; 28 trials). **g)** Bar graph summarizing the mean frequency pre-and 2 minutes post Θ stimulation for positive cases (***p<0.0001 pre-vs post stimulation). **h)** Bar graph summarizing the mean normalized frequency (% CTRL) for each 10 s bin during the baseline period and for the 4 min after Θ stimulation for positive cases (p<0.0001 pre-vs post stimulation). The blue bar describes mean frequency 8 min after the theta train.

All testing was counterbalanced by location of odors and treatment where applicable. Training and testing sessions were video recorded and later analyzed by an individual blind to group and treatment. A mouse was scored as exploring an odor when its nose was within 2 cm of, and directed towards, the odor hole. Odorants used in the above tests were as follows: (+)-Limonene (≥97% purity, Sigma-Aldrich); Cyclohexyl Ethyl Acetate (≥97%, International Flavors & Fragances Inc.); (+)-Citronellal (∼96%, Alfa Aesar); Octyl Aldehyde (∼99%, Acros Organics); Anisole (∼99%, Acros Organics); 1-Pentanol (∼99%, Acros Organics).

### >Partial unilateral silencing of field CA3 before behavioral testing

#### Stereotaxic surgery

Mice were anesthetized with ketamine and xylazine (100 and 10 mg/kg IP, respectively), and DREADD construct AAV8-CaMKIIa:HA-hM4Di-IRES-mCitrine (University of North Carolina Vector Core) or AAV8: pAAV-CaMKIIa-hM4D(Gi)-mCherry (Addgene, Watertown MA) was infused (2-3 min) into the target region using a 10-µl syringe with a 33GA metal needle (Hamilton, Reno, NV) as described^15^. The construct was infused unilaterally into CA3b using the following coordinates (AP, ML, and DV mm from suture) and quantities: (from bregma) −1.8, ±3.0, −2.4 (0.3 µl) and −2.9, ±2.87, −4.3 (0.5 µl). After surgery, mice were returned to their home cage for ≥ 4 weeks before behavioral testing to allow for virus expression.

#### Behavioral tests

Mice received an IP injection of vehicle or CNO (5 mg/kg; Tocris or National Institute of Mental Health) or vehicle (1% DMSO in saline), 30 min before the onset of cue sampling. Animals with Gi-DREADD injections were tested on ‘what’, ‘where’, or ‘when’ assays.

#### Histology

Animals were anesthetized after the conclusion of behavioral testing and decapitated; the forebrain was cryostat sectioned at 35 µm. Localization of the injection sites was assessed with fluorescence microscopy and by evaluating known projections from CA3 throughout the septo-temporal extent of the hippocampus. Three to four 370 µm thick hippocampal slices were prepared from a subgroup of animals and used for electrophysiological analyses of the extent to which CNO depressed synaptic transmission; these slices were subsequently sub-sectioned and examined to verify localization of fluorescent-DREADD markers. Animals that did not satisfy both injection placement and projection criteria were not included in the analysis of behavioral or electrophysiological data.

### Hippocampal Slice Electrophysiology

#### Recordings of pyramidal cell spikes from CA3

Hippocampal slices were prepared and transferred to an interface recording chamber (31 ± 1°C) as previously described^16,17^. Glass pipettes (R = 2-3 MΩ) containing 2M NaCl were placed in CA3a/b stratum pyramidale to record pyramidal cell (PC) spiking and evoked fEPSPs. Tests for evoked PC spiking used stimulating electrodes placed in CA3b/c and CA2/CA1 stratum radiatum (see Fig. 1a). The current intensity was adjusted so that dual stimulation produced a small composite fEPSP that typically was not accompanied by an antidromic spike. Extracellular recordings were fed through a Butterworth band pass filter (300-3000 Hz) and spiking analyzed off-line using custom-written computer code created with Python (v3.5), NumPy, and SciPy. There was considerable slice-to-slice variability in the amplitude of spikes and counting thresholds were adjusted accordingly (mean ± SEM: 100.1 ± 5.5 µV). The number of PC spikes was measured in 1 s bins and an average value calculated for each 10 s epoch. A mean frequency was measured over the 140 s baseline period (i.e. seven 10-s epochs spaced by 10 s) after which a 10-pulse Θ train was delivered and spike frequency similarly calculated for an additional 4 min. This stimulation protocol was repeated 2-3 times per slice, with 10 min between trains. Trials were dropped from the analysis if the coefficient of variation for the seven baseline epochs was greater than 45%; this occurred in 10.6% of the experiments. A mean baseline and post-stimulation PC spike frequency was calculated for all (2-3) trials for a given slice. For quantitative comparison between trials, within and across slices, the post-Θ data were normalized to their respective baseline frequency.

A separate set of experiments compared the effects of partially inhibiting the CA3 associational system, either through activation of Gi-DREADDs or by reducing ionotropic glutamate receptor function with 500 µM kynurenic acid (KYN), on spontaneous PC spiking and the amplitude of synaptic responses. A recording pipette was positioned in the CA3b PC layer to record spikes and the positive end of the fEPSP dipole. A single stimulating electrode was placed in CA1 stratum radiatum to activate Schaffer-commissural fibers terminating in the proximal apical dendrites of CA3. Pulses were applied at 3/min for at least 20 min prior to the application of KYN (40 min), CNO (10 µM, 30-40 min), or vehicle (30-40 min) to slices prepared from naive or Gi-DREADD expressing mice.

#### Drug Application

For hippocampal slice experiments, compounds were introduced to the bath via a second, independent perfusion line (6 ml/hr). CNO was made as a concentrated stock (1000X) in DMSO and then diluted to the bath concentration (10 µM) in ECS. The maximum DMSO concentration (≤0.01%) had no effect on baseline transmission. Kynurenic acid (Sigma Aldrich) was dissolved directly into ECS to achieve a final bath concentration of 500 µM.

## Results

We tested the hypotheses that: 1) field CA3, because of its massive collateral feedback projections^13^, operates as a regenerative network capable of prolonged, reverberating activity, and 2) partial silencing of the network will block the capacity to encode temporal order.

### Reverberating activity in the CA3 C/A system

Reverberation implies that activation of a recurrent network will increase mean firing rates by any given subpopulation of neurons for an extended period after the termination of the initiating input. We tested for such effects in hippocampal slices, preparations in which confounding effects of behaviorally driven stimulation are absent. Two electrodes were positioned to activate separate populations of C/A axons converging on field CA3b (**Fig 1a**). Single current pulses were adjusted to produce small (<400µV) fEPSPs in the target field and units were recorded from the pyramidal cell (PC) layer equidistant from the stimulation electrodes. Recordings were band pass filtered (300-3000 Hz) and an amplitude threshold (25-200 µV) used to detect individual spikes (**Fig 1b** and Methods). As expected for a network with dense recurrent connections, there was considerable spontaneous, arrhythmic spiking by CA3 PCs, something that is not found in field CA1 or in the granule cells of the dentate gyrus in hippocampal slices. Following a 2.5 min period of stable baseline firing, a physiologically relevant pattern of stimulation (a two second 10 pulse Θ-train [5 Hz]) was applied simultaneously to the stimulating electrodes and the subsequent 4 minutes of PC spiking recorded. From 23 slices, a total of 57 trials (i.e. 2-3 Θ trains separated by 10 min per slice) were accepted for analysis (Methods); post-stimulation firing rates for the multiple trials were averaged to obtain a single value for each slice. The mean spike frequency over the first 2 minutes after Θ stimulation was both highly correlated with spike frequency during the 2-minute baseline (R^2^=0.7783; **Fig 1c**) and elevated above the latter (baseline: 4.8 ± 0.6Hz; post-Θ: 5.7 ± 0.6 Hz; p=0.003) (**Fig 1d)**. Given the strong pre vs. post correlations, we normalized post-stimulation activity to its own baseline to allow for comparisons across trials and slices. Spike frequency was elevated immediately after the short Θ-train and remained so until the last 30 seconds of the 4-minute post-Θ session (**Fig 1e**). The strong correlations between pre-vs. post-stimulation periods confirm the stability of the slice preparations and indicate that factors regulating spiking rates continued operating after the short Θ train. Epileptiform discharges were not detected in traces or computerized analyses of spikes.

To gain a better perspective on the time course for the effects of Θ stimulation, we analyzed spike frequencies for trials in which positive effects (i.e., differences in pre-vs. post-spiking) were found. Nineteen (of 23) slices had at least one trial with an increase in PC spiking frequency after Θ. In total, there were 57 attempts to induce sustained activity; positive effects were obtained in 28 of these. There were no statistical differences in baseline activity between positive and negative cases or in the fEPSPs elicited by Θ stimulation. Positive trials were averaged to obtain a single value for each slice. As before, the mean spike frequency for the first 2 minutes after Θ stimulation was both highly correlated with spiking during the baseline across the 19 slices (R^2^=0.7677; **Fig 1f**) and elevated above the latter (baseline: 4.1 ± 0.2 Hz; post-Θ: 6.2 ± 0.7 Hz, p<0.0001) (**Fig 1g**). Post-Θ activity increased during the first 30 seconds to 50-60% above baseline and remained at that level for the remainder of the recording session (**Fig 1h**). This increase was not present in a 2-minute sample (mean of 7 time bins) collected 8 minutes after the 10-pulse Θ train (**Fig 1h**, last sample vs. original baseline p>0.05).

Complex systems with positive feedback are prone to catastrophic collapse or runaway activity with small changes in connection strength. We tested the point in hippocampal slices using the ionotropic glutamate receptor antagonist kynurenic acid and found that a concentration (500 µM) sufficient to depress C/A elicited fEPSPs by ∼20% caused a more than 70% decrease in the frequency of PC spiking (**Fig 2a**,**b**). A plot of the two measures over time of treatment emphasizes the extent to which firing frequency collapses with small changes in transmission (**Fig 2c**,**d**).

**Figure 2.**
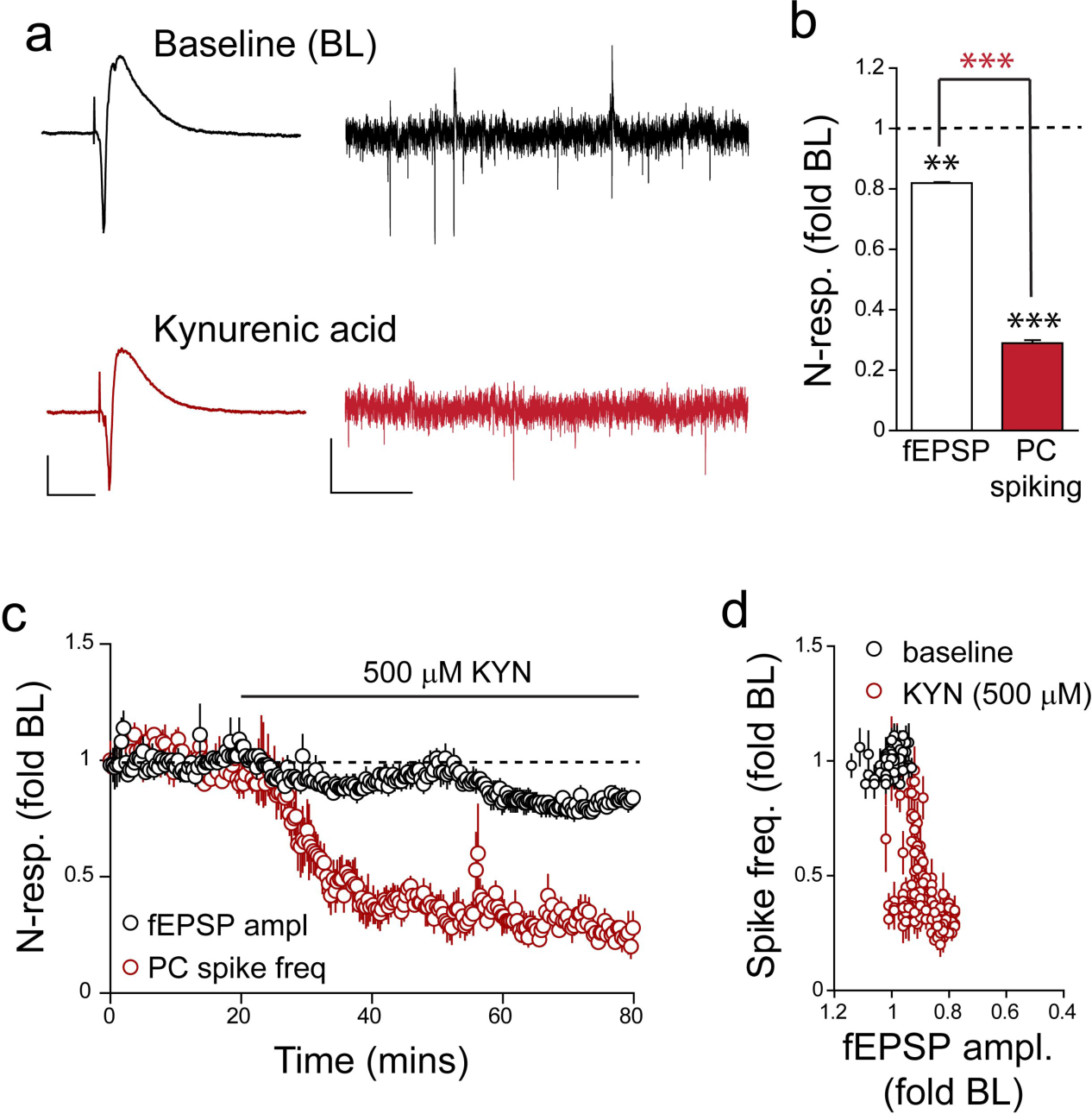
Autonomous spiking in CA3 depends on excitatory transmission. **a**) Representative CA3 fEPSPs (left) and traces of PC spiking (right) recorded in the absence (black) and presence (red) of kynurenic acid (KYN; 500 µM). The sharp negative deflection is the antidromic potential. Scale bars: fEPSP y=1mV, x=10ms; PC spiking y=100 μV, x=200ms. **b)** Bar graph summarizing the unequal effects of KYN upon fEPSP amplitude and PC spike frequency (n=4): **p=0.0014 for the effect on the fEPSP; ***p=0.0002 for the KYN effect on PC spiking; ***p=0.0007 for difference in KYN effect on fEPSP vs. spiking. **c)** Time course of KYN effects upon fEPSP amplitude and PC spiking. **d)** Graph of the reduction of PC firing vs. the decrease in fEPSP size during infusion of KYN. Panels c and d illustrate the catastrophic collapse in CA3 activity that occurs with small changes in transmission (n=4).

### Partial silencing of the CA3 C/A system selectively disrupts acquisition of temporal order

Field CA3 receives ∼50% of its pyramidal cell input from the contralateral side via the massive commissural system^13^. The sensitivity of the network to relatively small losses of excitatory transmission (see above results) strongly suggests that partial silencing of CA3 within one hemisphere will cause a bilateral reduction of recurrent firing. Note, however, that *rapid* throughput across the dentate gyrus-CA3-CA1 circuit would be minimally affected on the non-transfected side (**Fig 3a, right**). It follows from these arguments that unilateral suppression of CA3 can be used to evaluate the behavioral functions of the C/A network while leaving intact the process of identifying cues and their locations.

**Fig 3.**
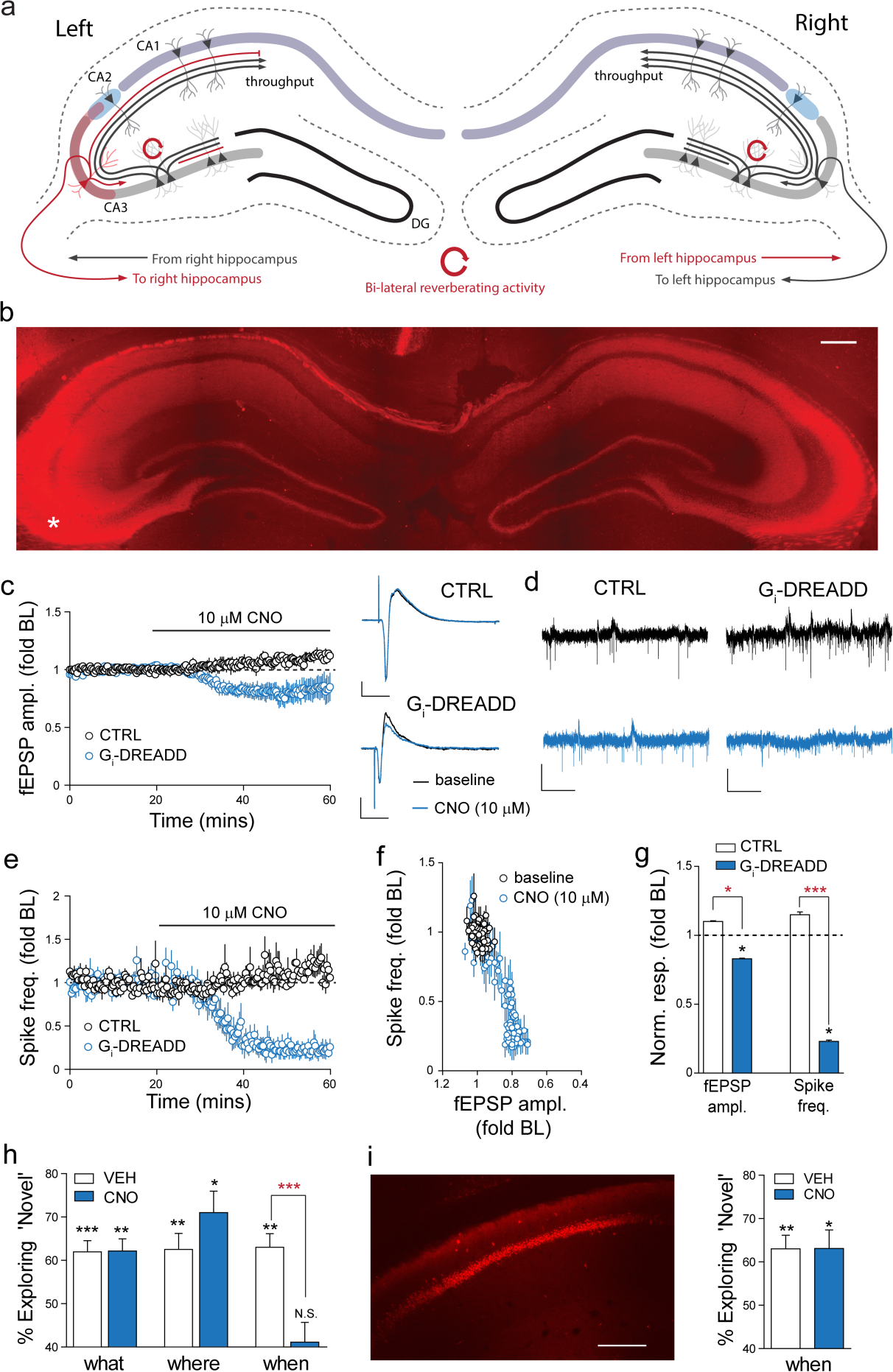
Unilateral silencing of a subpopulation of CA3 neurons selectively disrupts acquisition of temporal information. **a)** Design of experiments to selectively disrupt CA3 reverberation. The subfield receives afferents from entorhinal cortex and dentate gyrus and then monosynaptically innervates CA2 and CA1. This constitutes a rapid and potent throughput network. CA3 collaterals densely project within the ipsilateral subfield thereby forming an associational network capable of prolonged self-sustained activity. The systems present on each side are linked by a massive commissural projection that provides ∼ 50% of the pyramidal cell (PC) input to CA3 strata oriens and radiatum. Ipsilateral silencing of a portion of the CA3 system thus removes a significant portion of the feedback system on both sides of the hippocampus (red lines), a condition shown to cause catastrophic collapse of reverberation (red circles). **b)** Image shows the bilateral hippocampal distribution of mCherry following unilateral AAV-Gi-DREADD-mCherry transfection of field CA3a,b (asterisk, left marks injection placement; bar = 400 µm). **c)** Plot of Schaffer-commissural fEPSPs collected from stratum radiatum ipsilateral to, but outside, the transfected zone; CNO caused a modest reduction of these synaptic responses in transfected slices but had no effect on fEPSPs in non-transfected controls. Representative traces are shown on the right. Scale bars: y = 1 mV; x = 10 ms. **d)** Spiking by CA3 neurons prior to (black) and 40 min after (blue) CNO infusion in slices from control (CTRL, left) vs. Gi-DREADD transfected (right) hippocampi. Scale bars: y=100 µV, x=200 ms. **e)** Time course for CNO effects on mean spike frequency (± SEM) in slices from control vs. Gi-DREADD mice. **f)** Graph illustrating the catastrophic collapse in CA3 activity after bath application of CNO in CA3-DREADD mice (n=6. p=0.0003 fEPSP vs. spiking). **g)** Summary of the effects of CNO on fEPSP amplitude (CTRL: n=3, DREADD: n=8) and spiking (CTRL: n=3, DREADD: n=6) assessed at the conclusion of 40 min CNO infusion (*p≤0.05 paired t-test baseline vs CNO. Between group comparisons for control vs. DREADD: *p<0.05, ***p<0.0001, unpaired t-test). **h)** CA3-DREADD mice injected with CNO prior to sampling an odor sequence did not show the normal preference for an earlier vs. later odor during ‘when’ retention testing (‘N.S.’: p>0.05, paired t-test for n=8). Vehicle (VEH) injections did not disturb performance (**p=0.01). VEH vs. CNO: ***p<0.001 for % time sampling the earlier odor (unpaired t-test). CNO did not affect performance on the ‘what’ (n=7) or ‘where’ (n=6) tasks in CA3-DREADD mice (within group comparisons for time spent with novel cues or locations, respectively: *p=0.02; **p<0.01, ***p=0.0009, paired t-tests). **i)** Mice with unilateral transfection of CA1 (representative image, bar=400µm) and treated with CNO were not impaired in the ‘when’ task).

Accordingly, CA3a/b was unilaterally transfected with inhibitory Gi-DREADD construct in dorsal and ventral hippocampus (two injections). This resulted in construct expression by PCs across much of the septo-temporal extent of CA3a/b and dense transport to the ipsi-and contra-lateral targets of the C/A system (**Fig 3b**). We prepared hippocampal slices from CA3-DREADD mice after behavioral testing and confirmed that CNO produced a rapid reduction (<20%) in the size of CA3 fEPSPs elicited by stimulation of C/A projections in recordings collected outside the zone containing transfected cell bodies; CNO had no effect on C/A fEPSPs in slices from naïve control mice (i.e., not expressing the Gi-DREADD) (**Fig 3c**). The magnitude of the CNO effect on synaptic responses reflects the absence of transfection in a significant percentage of ipsi- and contra-lateral CA3 neurons that project to the recording site. While the DREADD agonist had a relatively modest effect on synaptic responses, it caused a profound reduction in cell firing recorded from the same electrode used to sample the positive end of the fEPSP dipole (**Fig 3d,e**). We estimated the relationship between % reductions in fEPSPs and CA3b spiking frequency by measuring the two variables at various intervals after onset of CNO infusion (**Fig 3f**) and obtained evidence for a catastrophic collapse of the network with a <20% decrease in the fEPSP (**Fig 3g**). We conclude from these analyses that the spatially restricted transfections had a relatively modest effect on throughput across field CA3 but produced a profound reduction in the capacity of the network to generate recurrent activity.

Unilateral CA3-DREADD mice given CNO prior to odor sampling had high retention scores in both the ‘what’ (p<0.002) and ‘where’ (p=0.02) paradigms described in methods (**Fig 3h**, what’ and ‘where’). The percent increase in exploration of the novel vs. familiar cue in the ‘what’ test was not significantly different between CNO- and vehicle-treatment groups; this was also the case for sampling of cues in new vs. previous locations in the ‘where’ problem. These findings confirm the prediction that depressing recurrent CA3 activity does not interfere with acquisition of information about the identity or location of multiple odor cues.

In striking contrast, unilateral CA3-DREADD mice given CNO did not acquire temporal order in the ‘when’ test. Vehicle treated transfected animals spent considerably more investigating the earlier odor (B>C) in the previously sampled A-D sequence in the retention trial (p=0.010) while those in the CNO group did not differentially explore the two cues (**Fig 3h** ‘when’). Unlike effects in CA3, unilateral Gi-DREADD transfection of field CA1 followed by CNO treatment did not disrupt the ability of mice to detect the order in which they had sampled the odors (VEH vs. CNO, p>0.05) (**Fig 3i**).

For unilateral CA3-DREADD mice, there were no evident differences between those receiving CNO vs. vehicle treatment on time spent investigating odors during the initial sampling period (p>0.05) or in exploration of the test chamber in the spatial paradigm (p>0.05). We conclude that partial inhibition of field CA3 and the associated disproportionate depression of cycling activity in the C/A network selectively interferes with acquisition of temporal order.

## DISCUSSION

A primary goal of the present experiments was to test the proposition that field CA3 is capable of generating responses of sufficient duration to bridge long gaps between serial cues, as routinely required by episodic memory^8^. This idea relates to the much discussed hypothesis that a sufficiently large population of interconnected neurons (cell assemblies^18^) can generate reverberating activity and thereby maintain a temporally extended representation for a brief input. Among other potential uses, systems of this kind are commonly assumed to provide a mechanism for working memory^19-22^. The cycling activity concept was developed from neuroanatomical considerations by Lorente de No (1933)^23,24^ and then used as a second postulate in Hebb’s influential theory of memory^18^. Reverberation also figured prominently in the first computer models of neural networks^25^ and has since been a topic of intense computational and theoretical work^22,26-28^. Perhaps the most convincing evidence for self-sustained activity in brain following a brief input comes from unit recording studies of primate frontal cortex during performance of a working memory task^20^. Firing in these cases generally lasts for a few seconds, as expected for working memory, but it is not entirely clear that the effects are due to the type of reverberation envisioned by Hebb and other theorists. Our experiments found that the unusually dense interconnections within CA3 are capable of maintaining autonomous activity for much longer periods: a two second input initiated firing that in many cases persisted for minutes. Given that recordings were collected in an isolated preparation without peripheral input, and were sensitive to modest changes in the network, it can be concluded the prolonged activity was intrinsically generated.

These results suggested a means for avoiding a major difficulty facing attempts to use experimental manipulations to investigate the contributions of the C/A system to memory. Bilateral silencing of a substantial portion of CA3 will interrupt the primary internal circuit of the hippocampus (DG-CA3-CA1) and thus block most forms of processing performed by the structure. Unilateral silencing of a fraction of CA3 cells will eliminate a significant percentage of the bilateral C/A system, and thus reduce cycling between the two hemispheres, while leaving fast throughput intact on the contralateral side.

To investigate behavioral consequences, we used protocols that as with human episodic memory did not involve prior training, repetition, or rewards. These assays sampled acquisition of three essential features of an episode: the identity, location, and temporal order of multiple cues (respectively, ‘what, ‘where’, and ‘when’). Unilateral chemogenetic silencing of a subpopulation of CA3 pyramidal cells blocked retention of the temporal order in which cues were sampled but did not detectably affect acquisition of ‘what’ and ‘where’ information. This latter result confirms that opposite side processing was intact. Given that the animals remembered the individual cues and their locations, we conclude that the CA3 system attached temporal information to the representations of the cues.

It seems unlikely that the recursive network maintains firing associated with each cue, and keeps these segregated, throughout the sampling period and the delay before retention tests. A more plausible explanation is that the prolonged spiking produced by the system gradually produces a transient potentiation of synapses between CA3 neurons participating in the cycling initiated by an odor, an effect that could persist for the duration of the sessions. The steady decay of enhanced synaptic responses elicited by the odors would be proportional to the order in which the cues had occurred, thereby providing a relative novelty basis for distinguishing between them in the retention trial. An alternative hypothesis for the results in the temporal order test is that prolonged firing in CA3 results in linkages between the separate populations of neurons activated by successive members of the cue sequence. In this scenario, presentation of two odors together in the retention test would result in a prompted (stronger) response to the later cue in the original sequence and thus greater perceived novelty of the earlier one during the retention trial.

In summary, the present findings indicate that a fully intact bilateral CA3 system is required for a critical temporal component of episodic memory but not for acquiring two other essential elements. Physiological analyses showed that the singular CA3 feedback system has a capacity for self-sustained activity lasting for unprecedented periods and thus is sufficient to provide information across long intervals between cues. We propose that this capacity is critical to performance in episodic ‘when’ tests.

## Acknowledgements

The authors thank Nima R. Hadidi and Johnny Q. Nguyen for assistance with behavioral studies. This work was supported by Office of Naval Research Grant N00014182114, NIMH grant MH082042 and NIH grant HD89491., B.M.C was supported by the NIH National Center for Research Resources and National Center for Advancing Translational Sciences UL1 TR001414. C.D.C. was supported by NIH T32 AG00096-34.

